# An intrinsically disordered kinetochore protein coordinates mechanical regulation of chromosome segregation by dynein

**DOI:** 10.1101/2023.05.07.539709

**Authors:** Jessica M. McGory, Dylan M. Barcelos, Vikash Verma, Thomas J. Maresca

## Abstract

Kinetochores connect chromosomes and spindle microtubules to maintain genomic integrity through cell division. Crosstalk between the minus-end directed motor dynein and kinetochore-microtubule attachment factors promotes accurate chromosome segregation through a poorly understood pathway. Here we identify a physical linkage between the intrinsically disordered protein Spc105 (KNL1 orthologue) and dynein using an optogenetic oligomerization assay. Core pools of the checkpoint protein BubR1 and the adaptor complex RZZ mediate the connection of Spc105 to dynein. Furthermore, a minimal segment of Spc105 that contains regions with a propensity to multimerize and binding motifs for Bub1 and BubR1 is sufficient to functionally link Spc105 to RZZ and dynein. Deletion of the minimal region from Spc105 reduces recruitment of its binding partners to bioriented kinetochores and causes chromosome mis-segregation. Restoration of normal chromosome segregation and localization of BubR1 and RZZ requires both protein binding motifs and higher-order oligomerization of Spc105. Together, our results reveal that higher-order multimerization of Spc105 is required to recruit a core pool of RZZ that modulates microtubule attachment stability to promote accurate chromosome segregation.

## INTRODUCTION

Kinetochores serve as the linkage between the centromeric region of chromosomes and the spindle microtubules that facilitate chromosome segregation to daughter cells. The affinity of kinetochores for spindle microtubules is carefully modulated to support proper chromosome segregation. Without tight regulation, defects in kinetochore-microtubule (kMT) attachments arise and lead to segregation errors, ultimately compromising genomic integrity and resulting in aneuploidies. Modulation of this attachment is thought to be governed by two pathways-a phospho-regulation pathway and a mechanical or steric regulatory pathway. While phospho-regulation of kMT affinity has been extensively studied (Barbosa et al., 2022; Broad and DeLuca, 2020; Maresca and Salmon, 2010; Santaguida and Musacchio, 2009), a more recently described “crosstalk” pathway (Cheerambathur et al., 2013) involving mechano-regulation of kMT attachment factors by the motor protein dynein is poorly understood. Interestingly, the core kMT attachment sites that are regulated by dynein are located in the outer kinetochore in a milieu of the highly enriched intrinsically disordered protein KNL1 (Spc105 in *Drosophila*) (**Fig. 1 A**).

**Figure 1.**
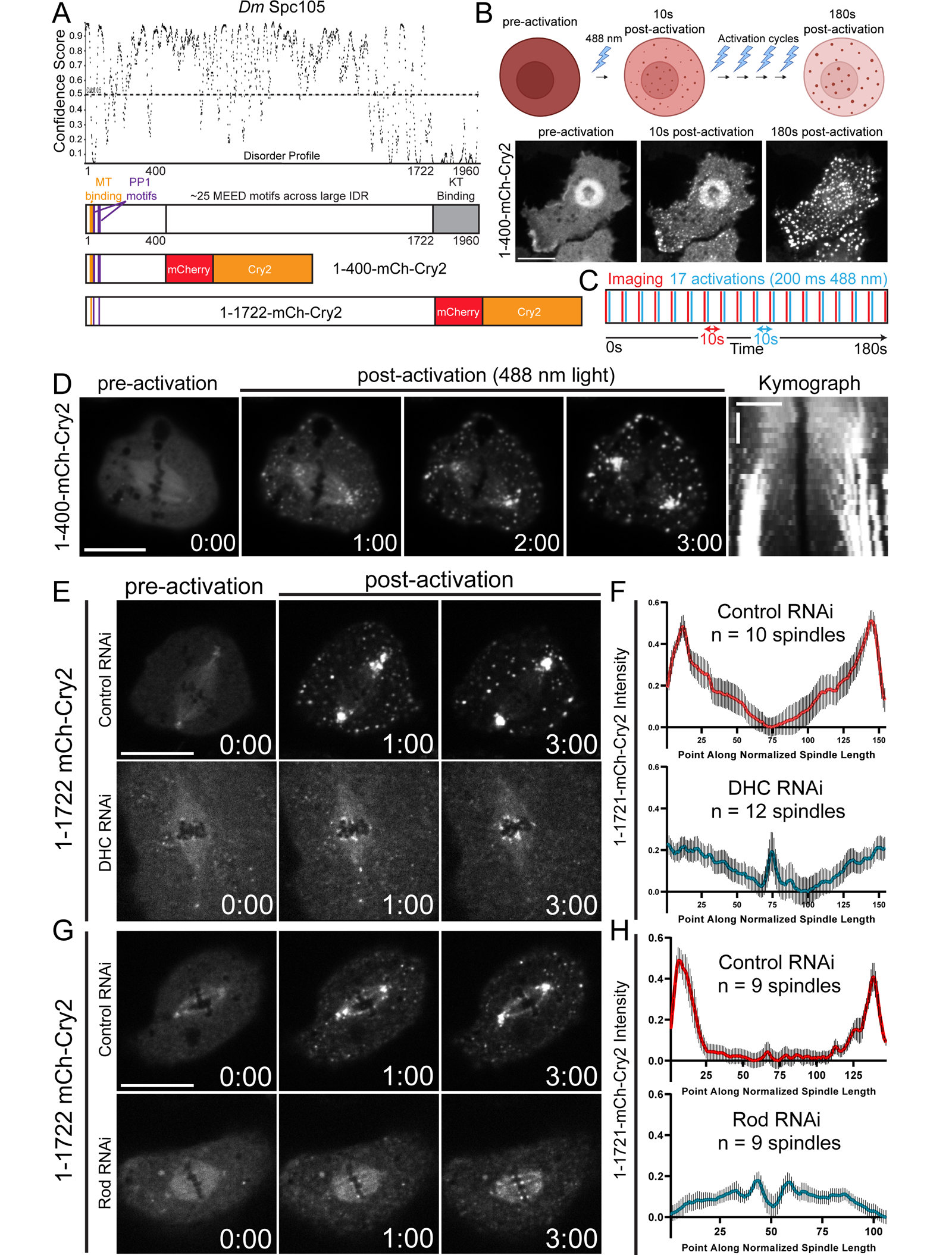
Oligomerized Spc105 exhibits dynein- and RZZ-dependent poleward movement. (A) Disorder plot of Spc105 and schematics of full length Spc105 and truncations fused to mCherry-Cry2. (B) Schematic of the photo-oligomerization assay with a representative example of its application in living cells. (C) Schematic of the activation/imaging protocol applied to visualize oligomers in mitotic cells. (D) Representative example of photo-oligomerized 1-400-mCherry-Cry2 streaming poleward on a mitotic spindle with accompanying kymograph. (E) Representative control and DHC-depleted cells expressing 1-1722-mCherry-Cry2 and subjected to the activation/imaging protocol. (F) Quantification of the distribution of 1-1722 oligomers across mitotic spindles in control (red) and DHC-depleted (cyan) cells 3-minutes post activation (Control, n = 10 spindles; DHC RNAi, n = 12 spindles). (E) Representative control and Rod-depleted cells expressing 1-1722-mCherry-Cry2 and subjected to the activation/imaging protocol. (F) Quantification of the distribution of 1-1722 oligomers across mitotic spindles in control (red) and Rod-depleted (cyan) cells 3-minutes post activation (Control, n = 9 spindles; Rod RNAi, n = 9 spindles). Error bars are SEM. Scale bars, 10 μm; kymograph; 5 μm (horizontal) and 1 min (vertical). Elapsed time is min:sec.

## RESULTS AND DISCUSSION

The N-terminal region of KNL1 has been implicated in Aurora B kinase-mediated phospho-regulation of kMT attachment stability in human cells (Caldas et al., 2013); however, we previously reported that deletion of the first 400 amino acids (Δ400) of Spc105 caused hyper-stable kMT attachments without evidently affecting Aurora B kinase activity in *Drosophila melanogaster* S2 cells (Audett et al., 2022). The N-terminus of KNL1 orthologues have conserved PP1 and MT binding motifs (**Fig. 1 A**), however, restoration of these activities to the Spc105Δ400 mutant did not restore normal kMT attachment stability. Thus, we hypothesized that the N-terminal 400 aa of Spc105 recruits additional activities that weaken kMT interactions independent of Aurora B kinase. In our prior work (Audett et al., 2022), the first 200 amino acids of Spc105 was fused to a minimal region of the *Arabidopsis thaliana* protein CRY2 that oligomerizes in response to blue light (Bugaj et al., 2013). The Spc105-Cry2 oligomerization assay recapitulates - in the cytosol - the complex multivalent nature of the kinetochore where there are greater than 100 of copies of Spc105. This optogenetic assay bypasses many technical challenges with studying kinetochore protein functions in the context of intact kinetochores under physiological conditions that are challenging to reproduce *in vitro*. Thus, we deployed the Cry2-based optogenetic oligomerization assay to gain insights into kMT attachment-regulatory activities based in the first 400 aa of Spc105 (**Fig. 1 A**).

Spc105-1-400-mCherry-Cry2-expressing cells assembled cytoplasmic oligomers upon exposure to 488 nm laser light (**Fig. 1 B**). An activation/imaging protocol was applied to mitotic cells to examine the behavior of the 1-400 oligomers with relationship to the mitotic spindle (**Fig. 1 C**). Strikingly, 1-400 oligomers not only associated with the mitotic spindle, but also moved poleward and accumulated at spindle poles following photo-activation (**Fig. 1 D, Video 1**). Further substantiating the physiological relevance, an mCherry-Cry2-tagged version of Spc105 containing the entire intrinsically disordered region (IDR) but lacking the C-terminal kinetochore targeting domain (1-1722-mCherry-Cry2) also photo-oligomerized and streamed poleward on the spindle (**Fig. 1 A, E**). The poleward movement of the Spc105-mCherry-Cry2 oligomers implicated the minus-end directed motor dynein, which is known to be targeted to the kinetochore by the Rod-Zw10-Zwilch (RZZ) complex (Gama et al., 2017; Mosalaganti et al., 2017). Upon depletion of the dynein heavy chain (DHC) by RNAi (**Fig. S1 A-C**), Spc105 oligomers did not accumulate at spindle poles, but rather became enriched near the kinetochore region (**Fig. 1 E, F and Video 2**). Since RZZ recruits the checkpoint proteins Mad1 and Mad2 (Caldas et al., 2015; Kops and Gassmann, 2020), which are stripped from kinetochores in a dynein-mediated manner that is remarkably similar to the behavior of Spc105 oligomers (Basto et al., 2004; Cane et al., 2013; Howell et al., 2000; Howell et al., 2001), we next assessed the motility of the 1-1722 oligomers in Rod-depleted cells (**Fig. S1 D-F**). Comparable to the DHC RNAi condition, 1-1722 oligomers in Rod-depleted cells did not move poleward and accumulated near kinetochores rather than at spindle poles (**Fig. 1 G, H and Video 3**). Thus, the minus-end directed movement and polar accumulation of Spc105 oligomers required both dynein and its adaptor - the RZZ complex.

We next examined the 1-400 region of Spc105 for sequence features that could explain the molecular mechanism for the functional linkage of Spc105 to RZZ and dynein revealed by the oligomerization assays. T-COFFEE sequence alignment (Notredame et al., 2000) of the Spc105 and human KNL1 amino acid sequences identified two putative KI motifs (KI1 and KI2) in the N-terminal region of Spc105 between amino acids amino acids 279-349 (**Fig. 2 A**). In human KNL1, KI motifs bind directly to the N-terminal TPR motifs of the Bub checkpoint proteins with KI1 and KI2 exhibiting preferential binding for Bub1 and BubR1 respectively (Kiyomitsu et al., 2011; Kiyomitsu et al., 2007; Krenn et al., 2012). We investigated this potential relationship for Spc105 using the oligomerization assay by fusing 266-384 to mCherry-Cry2. In support of the 266-384 region containing functional KI motifs, both Bub1-EGFP and BubR1-EGFP were enriched in the 266-384 oligomers with BubR1 exhibiting ∼2-fold higher enrichment in the clusters than Bub1 (**Fig. 2 C, D, G, J, K, L**). Importantly, the binding was specific as the EGFP tag alone was not enriched in the 266-384 oligomers (**Fig. 2 B**). We additionally noted that N-terminally tagged Bub1 was not recruited to the 266-384 oligomers, (**Fig. 2 C**) which may be due to the N-terminal EGFP tag interfering with binding of the TPR to the KI1 motif. Furthermore, we were able to rule out that enrichment of Bub1 and BubR1 in 266-384 clusters was mediated by the checkpoint protein Bub3, which binds to and recruits subpopulations of Bub1 and BubR1 to intact kinetochores (Krenn et al., 2012; Larsen et al., 2007; Overlack et al., 2015; Primorac et al., 2013; Taylor et al., 1998), because the KI region does not contain any Bub3 binding motifs and we did not observe enrichment of Bub3 in the 266-384 oligomers (personal observation). Altogether, the data supports the conclusion that Spc105 contains functional KI motifs although we postulate that the KI2 motif may be more “active” since it is better conserved than the KI1 motif sequence and BubR1 was more robustly recruited to 266-384 oligomers than Bub1. Since the N-terminus of KNL1 and Bub1 contribute to localization of the RZZ complex at human kinetochores (Caldas et al., 2015; Varma et al., 2013; Zhang et al., 2019; Zhang et al., 2015), we next investigated whether RZZ components were recruited to 266-384 oligomers. Consistent with prior findings, Zw10-EGFP and superfolder GFP (sfGFP)-Rod were similarly enriched in the 266-384 drops (**Fig. 2 C, E, F, H-L**). Interestingly, 266-384 clusters that recruited sfGFP-Rod moved poleward on mitotic spindles (**Fig. 2 M and Video 4**). Thus, oligomerization of a minimal KI region (266-384) was sufficient to recruit the molecular machinery linking Spc105 to dynein and to support the minus-end directed motility observed for oligomerized 1-400 and 1-1722 Spc105 fragments.

**Figure 2.**
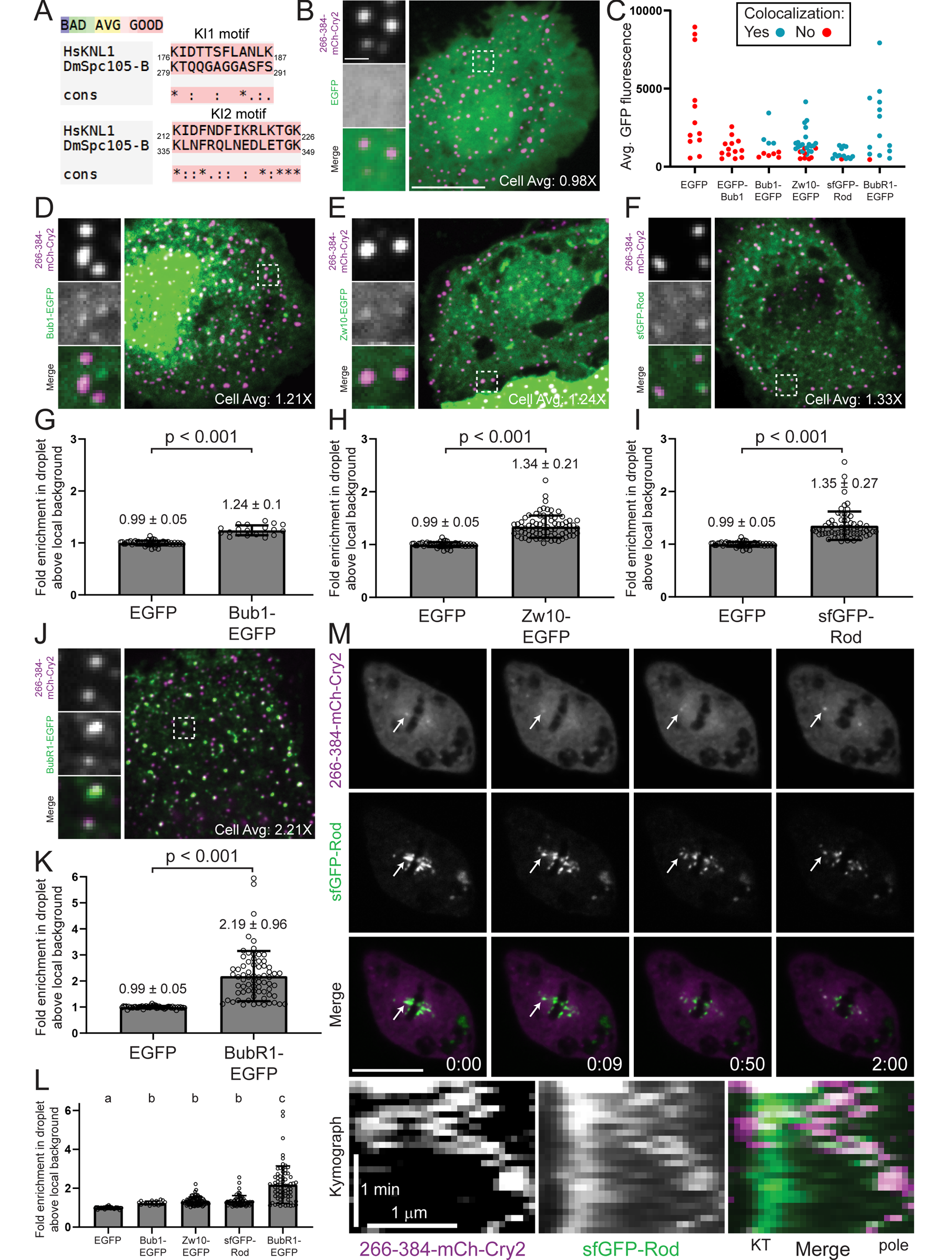
Oligomerization of a minimal KI region of Spc105 recruits binding partners and is sufficient for poleward movement. (A) T-COFFEE alignments of human (Hs) KNL1 and Spc105 identify putative KI motifs. (B, D, E, F, J) Representative confocal Z-planes of cells co-expressing the indicated EGFP-tagged protein (green) and oligomerized clusters of the minimal KI motif-containing region (266-384) fused to mCherry-Cry2 (magenta). (C) Co-localization of the indicated EGFP-tagged proteins with 266-384-mCherry-Cry2 clusters in cells over a range of EGFP expression. Co-localization was never observed in cells with a broad range of EGFP (control) or EGFP-Bub1 expression levels. With the exception of the EGFP control (to establish a baseline measurement for the assay), only cells exhibiting co-localization were used to measure the fold-enrichment for each binding partner. (G, H, I, K) Quantification of the fold enrichment of the signal of the indicated EGFP-tagged protein relative to the local background signal (EGFP; n = 53 drops from 10 cells, Bub1-EGFP; n = 20 drops from 4 cells, Zw10-EGFP; n = 80 drops from 16 cells, sfGFP-Rod; n = 65 drops from 13 cells, BubR1-EGFP; n = 70 drops from 14 cells). (L) Measurements from G, H, I, and K shown on the same plot. In all plots, each binding partner is compared to the same control EGFP data set. (M) Representative confocal time-lapse of sfGFP-Rod (green) co-localized with a 266-384-mCherry-Cry2 cluster (magenta) moving poleward (arrow) with accompanying kymograph. Boxed regions are highlighted in the insets. The average fold enrichment for the representative cell is indicated in each image. Error bars are SD. Scale bars, 10 μm; 1 μm (insets), and 1 μm (horizontal); 1 min (vertical) in kymograph. Letters above each plot column indicate significant differences (p < 0.05) determined by a randomization method for all pairwise combinations. Same letters indicate that the difference is not significant (p > 0.05).

While region 266-384 was sufficient to enrich BubR1 and RZZ in cytosolic clusters, we next assessed if the region was necessary for recruitment of these proteins to intact kinetochores by deleting aa 266-384 from full-length Spc105. Bioriented kinetochores in the Δ266-384 deletion mutant exhibited approximately 45% and 60% reductions in BubR1 and Rod levels, respectively, compared to cells expressing wildtype (WT) Spc105 (**Fig. 3 A-D**). Since we postulate that BubR1 binds directly to Spc105 via the KI2 motif, we depleted BubR1 via RNAi to determine if RZZ recruitment to bioriented kinetochores was downstream of BubR1 (**Fig. S1 G, H**). Depletion of BubR1 from WT cells resulted in a comparable reduction in Rod levels at bioriented kinetochores as was observed in the Δ266-384 deletion mutant (**Fig. 3 D-F**), suggesting that BubR1 bridges the KI2 motif of Spc105 and RZZ. Effects on chromosome segregation were next evaluated via live and fixed cell analyses to determine if the 266-384 region was necessary for normal kinetochore function. Consistent with prior work in human cells expressing a mutant version of KNL1 lacking the KI motifs (Kiyomitsu et al., 2011), there was a significant increase in the frequency of lagging chromosomes during anaphase in cells expressing Spc105 Δ266-384-EGFP compared to control cells (**Fig. 3 G, H and Fig. S2 A**). Upon evaluation of the kMT attachment states of the lagging chromosome, we measured a ∼6-fold increase in the frequency of merotelic attachments, an erroneous kMT attachment in which a single kinetochore is attached to microtubules oriented towards opposite spindle poles (**Fig. 3 I**). In addition, the 266-384 deletion mutant exhibited a ∼6-fold increase relative to control cells in the frequency of cut chromatin in telophase/cytokinesis; most likely due to the persistence of merotelic attachments through anaphase as the cleavage furrow pinched lagging chromosomes (**Fig. 3 G, I**). In live-cell confocal microscopy, merotelic attachments often became deformed into characteristic bilobed/kidney-shaped structures (Cimini et al., 2001) as they were pulled in opposing directions during anaphase (**Fig. 3 J and Video 5**). The abundance of persistently stabilized merotelic attachments in the 266-384 deletion mutant supported the conclusion that loss of the KI motifs contributed to the hyper-stable kMT attachments we observed in the Δ1-400 Spc105 mutant (Audett et al., 2022).

**Figure 3.**
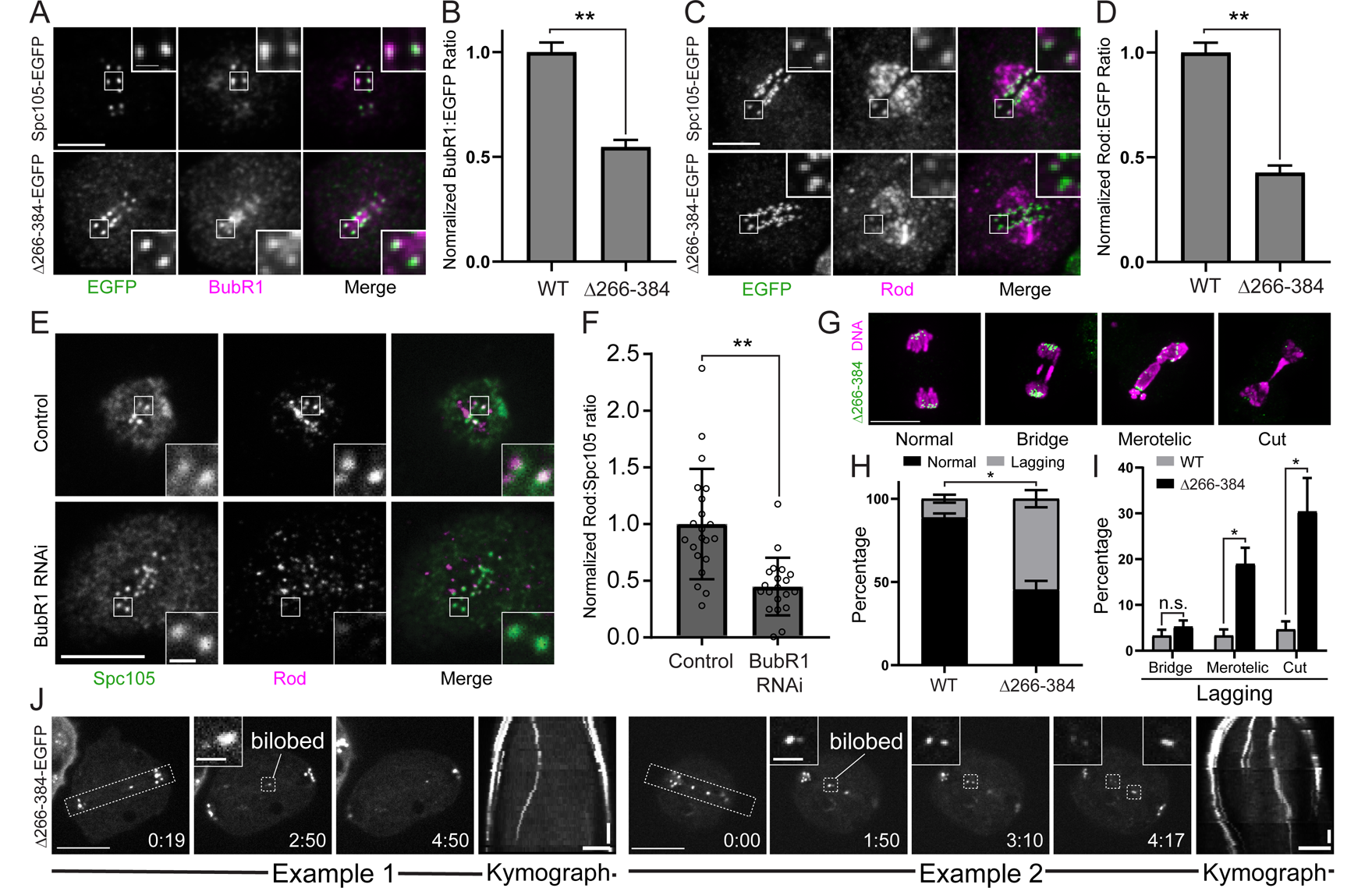
Dissecting the molecular linkage between Spc105 and dynein and its functional contribution to accurate chromosome segregation. (A) Representative images of BubR1 (magenta) in cells expressing WT Spc105 or the Δ266-384-Spc105 deletion mutant (green). (B) Quantification of BubR1 intensity relative to EGFP signal in WT and Δ266-384-expressing cells (WT; n = 140 kinetochores from 25 cells and 2 replicates, Δ266-384; n = 135 kinetochores from 24 cells and 2 replicates). BubR1 was often enriched at the inner centromere in cells expressing the mutant. (C) Representative images of Rod (magenta) in cells expressing WT Spc105 or the deletion mutant (green). (D) Quantification of Rod intensity relative to EGFP signal in WT and Δ266-384-expressing cells (WT; n = 142 kinetochores from 25 cells and 2 replicates, Δ266-384; n = 140 kinetochores from 25 cells and 2 replicates). (E) Representative confocal Z-planes of control and BubR1 dsRNA-treated fixed cells stained for Spc105 (green) and Rod (magenta). (F) Quantification of Rod:Spc105 intensity ratios in control and BubR1-depleted cells (control; n = 105 kinetochores from 21 cells and 2 replicates, BubR1 RNAi; n = 102 kinetochores from 20 cells and 2 replicates). The data points in the plot represent cell averages from measuring ∼5 kinetochores/cell. (G) Examples of the chromosome segregation categories quantified in I (Δ266-384 is green, DNA is magenta). (H) Percentage of anaphase/telophase cells with at least one lagging chromosome in WT- and Δ266-384-expressing cells (n = 150 cells per condition from 3 replicates). (I) Percentage of WT- and Δ266-384-expressing cells exhibiting each of the lagging chromosome phenotypes. (J) Representative examples of Δ266-384-expressing cells with persistent merotelic attachments in anaphase. Kymographs of the boxed regions are shown. Reported p-values were generated by a randomization method in B, D, and F or two-tailed t-test in H and I: * p-value < 0.05, ** p-value < 0.001. Boxed regions are shown in the insets. All error bars are SEM except for SD in F. Scale bars: 10 μm (E, G, J); 5 μm (A, C), 1μm (insets) (A, C, E, J); 5 μm (horizontal) and 1 min (vertical) in kymographs. Elapsed time is min:sec.

While implementing the activation/imaging protocol, we noted that oligomerization occurred more readily in cells expressing 1-400- and 1-1722-mCherry-Cry2 than in control cells with comparable levels of mCherry-Cry2 expression (**Fig. 4 A and Video 6**). Accordingly, the mean expression levels (determined by measuring the average mCherry fluorescence intensities of the cytoplasm prior to photo-activation) that supported Cry2-based clustering was statistically significantly different for each protein with mCherry-Cry2 requiring the highest expression, 266-384 being intermediate, and 1-1722 being the lowest (**Fig. 4 B**). The Cry2-based oligomerization assay was previously adapted to study the ability of “sticky” IDRs, which we believe aptly describes Spc105, to form so-called optoDroplets in living cells (Shin et al., 2017). Interestingly, oligomerized Spc105 exhibited droplet-like properties as 266-384 clusters could be observed fusing in the cytosol (**Fig. 4 C and Video 7**). Furthermore, introduction of the alcohol 1,6-hexanediole (1,6-HD), which disrupts weak hydrophobic interactions, rapidly disassembled the 266-384 clusters (**Fig. 4 D and Video 8**).

**Figure 4.**
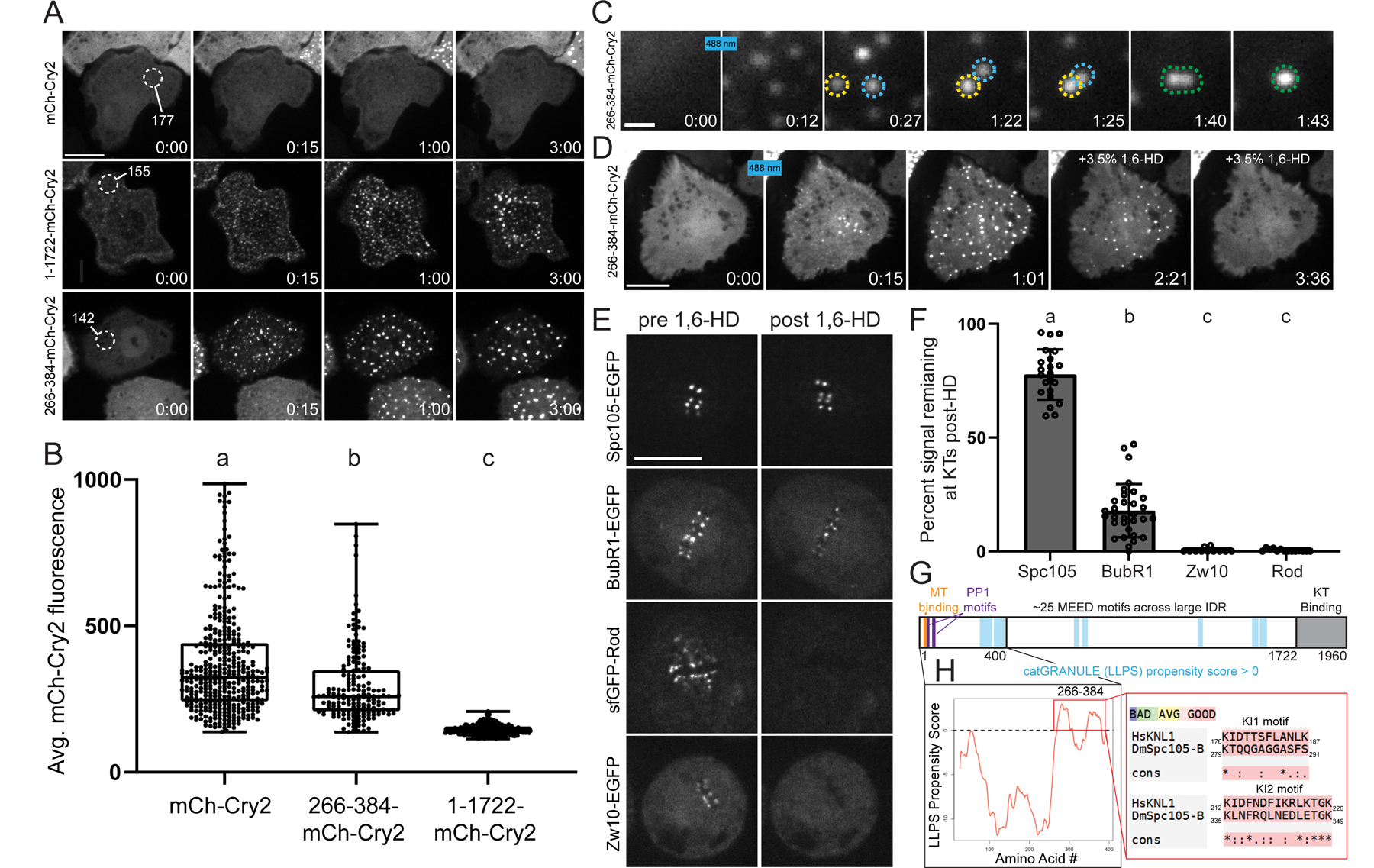
Characterization of cytosolic Spc105 oligomers and how hydrophobic interactions contribute to the Spc105-dynein linkage at intact kinetochores. (A) Representative time-lapse confocal imaging of cells with comparable expression of the mCherry-Cry2 tag alone or tagged Spc105 truncations subjected to the same activation protocol. (B) Box and whisker plots of the average pre-activation mCherry fluorescence of all the cells that formed clusters in the activation/imaging protocol. (C) Representative confocal time-lapse of 266-384-mCherry-Cry2 clusters (yellow and blue circles) fusing (green circle). (D) Representative confocal time-lapse of optogenetic clustering in a 266-384-mCherry-Cry2-expressing cell being rapidly reversed upon the addition of 3.5% 1,6-HD. (E) Still frames from spinning disc confocal time-lapses of cells expressing the indicated EGFP-tagged protein prior to treatment and three minutes after introducing 3.5% 1,6-HD into the imaging chambers. (F) Quantification of the EGFP signal for each of the indicated proteins remaining at kinetochores three minutes after introducing 3.5% 1,6-HD into the imaging chamber (Spc105; n = 22 cells, BubR1; n = 31 cells, Zw10; n = 11 cells; Rod; n = 14 cells). (G) Schematic of *D. melanogaster* Spc105-B highlighting the N-terminal MT-(orange) and PP1-binding (magenta) motifs and the seven regions across Spc105 with a catGRANULE LLPS propensity score above 0 (blue). (H) catGRANULE plot of the first 400 aa of Spc105-B with region 266-384 highlighted in the inset showing T-COFFEE alignments of the overlapping KI motifs (Hs*: Homo sapiens;* Dm*: D. melanogaster*). Whiskers indicate min-max, box is 25^th^-75^th^ percentile, and bar is median in B. Error bars are SD in F. Scale bars, 10 μm. Letters above each plot column indicate significant differences (p < 0.05) determined by a randomization method for all pairwise combinations. Same letters indicate that the difference is not significant (p > 0.05). Scale bars, 10 μm (A, D, E); 1 μm (C). Displayed times are min:sec.

A hydrophobic region in the first ∼240 amino acids of *C. elegans* KNL-1 mediates higher-order oligomerization of KNL-1 *in vitro* (Kern et al., 2015). Consistent with this phenomenon, 1,6-HD reduced the extent to which Spc105 (1-400) bound MTs *in vitro*, which we interpreted to be a consequence of disrupting Spc105 homo-oligomerization via hydrophobic interactions (Audett et al., 2022). While 1,6-HD impacted Spc105 function *in vitro*, we next assessed the effects of 1,6-HD on the phospho-(Aurora B kinase-mediated) and mechano-(RZZ-mediated) regulatory pathways that modulate kMT attachment stability in living cells. Interestingly, addition of 1,6-HD did not impact the phospho-regulatory pathway in fly cells as levels of active Aurora B kinase at the inner centromere relative to Spc105 were indistinguishable between 1,6-HD-treated and control cells (**Fig. S2 B, C**). Conversely, the core pool of the RZZ complex at bioriented kinetochores was dramatically impacted by the treatment as Rod and Zw10 rapidly and completely dissociated from kinetochores upon introduction of 3.5% 1,6-HD. The loss of the core pool of RZZ in the presence of 1,6-HD was not a consequence of significant Spc105 mis-localization and BubR1 levels were reduced but not to the same extent as Zw10 or Rod (**Fig. 4 E, F, Fig. S2 D and Video 9**). Since we could not ignore the facts that: 1) weak hydrophobic interactions between IDRs are implicated in a higher-order oligomerization phenomenon known as liquid-liquid phase separation (LLPS) and 2) 266-384 oligomers exhibited LLPS-like properties, we further examined the sequence features of Spc105 using the *cat*GRANULE algorithm (Bolognesi et al., 2016) to assess the propensity of Spc105 to phase separate. The algorithm identified seven stretches (between 8-40 aa) in Spc105 that may have a propensity to phase separate (**Fig. 4 G**). Consistent with the observed properties of 266-384 oligomers (**Fig. 4 C, D**), the two longest (40 and 35 aa) and highest scoring regions of Spc105 mapped between aa 266-384 and largely overlapped with the KI motifs (**Fig. 4H**).

Since the LLPS regions overlapped with the KI motifs, it was not possible to discern whether the Δ266-384 mutant phenotype was due to loss of the protein binding motifs, reduced ability to assemble higher-order oligomers, or both. To overcome this challenge, we took advantage of the KI motifs in human KNL1, which do not overlap with predicted LLPS regions in the N-terminus of KNL1 (**Fig. S3 A**). We first replaced the Spc105 KI region with the human KI motifs (*Hs*KI) via insertion into the Δ266-384 deletion to test if it was sufficient to recruit BubR1 and Rod. Upon inclusion of the *Hs*KI motifs, BubR1 and Rod localization was not restored at bioriented kinetochores as they were reduced ∼35% and ∼60% respectively compared to the control cells (**Fig. 5 A-D**). We next replicated the chimera experiment with the addition of the IDR of human FUS, which assembles into higher-order oligomers via phase-separation. The properties of FUS have been thoroughly characterized in recent years (Patel et al., 2015; Shin et al., 2017), and FUS has no known role at the kinetochore, making it an ideal candidate to assess the contribution of higher-order oligomerization to BubR1 and Rod localization. The FUS-*Hs*KI-expressing cells exhibited levels of BubR1 and Rod recruitment that were higher than the *Hs*KI alone, and not statistically significantly different from the levels in WT Spc105-expressing cells (**Fig. 5 E-H**), indicating a near-full rescue of protein localization. We next tested the contribution of these regions to the lagging chromosome phenotype to determine the functionality of the chimeras. Cells expressing the Δ266-384 deletion and the *HsKI* chimera both exhibited a significant increase in lagging chromosomes compared to control cells, while addition of both the *Hs*KI motifs and the FUS IDR alleviated the segregation defects (**Fig. 5 I**). Similarly, the frequency of merotelic or “cut” chromatin bridges was elevated in cells expressing the Δ266-384 deletion and the *Hs*KI chimera but comparable between WT- and FUS-*HsKI*-expressing cells (**Fig. 5 J**).

**Figure 5.**
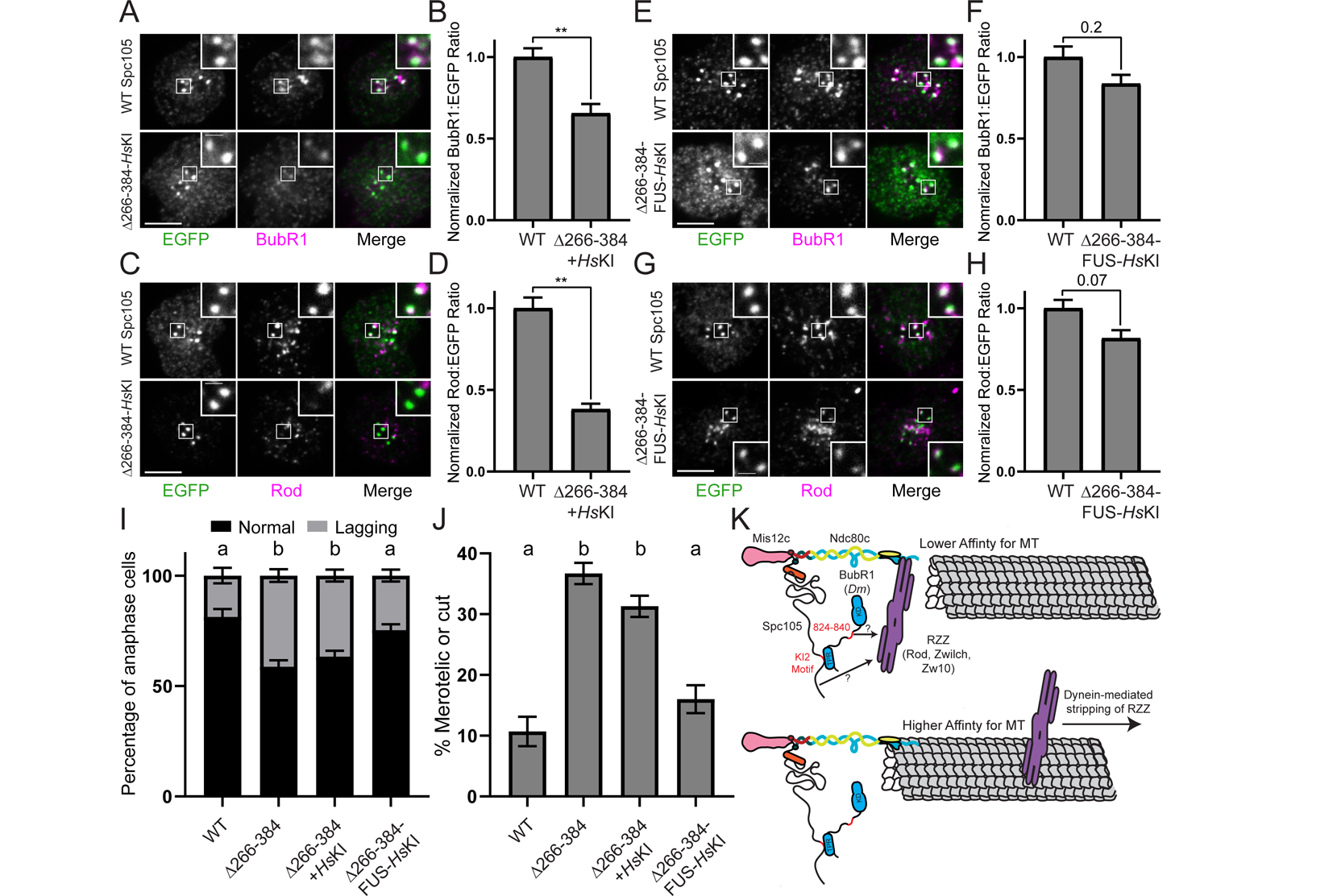
Localization of dynein targeting factors and accurate chromosome segregation require higher order multimerization and protein binding motifs in Spc105. (A) Representative images of BubR1 (magenta) in cells expressing WT Spc105 or the *Hs*KI chimera (green). (B) Quantification of BubR1 intensity relative to EGFP signal in WT and *Hs*KI-expressing cells (WT; n = 150 kinetochores from 30 cells and 3 replicates, *Hs*KI; n = 145 kinetochores from 29 cells and 3 replicates). (C) Representative images of Rod (magenta) in cells expressing WT Spc105 or the *Hs*KI chimera (green). (D) Quantification of Rod intensity relative to EGFP signal in WT and *Hs*KI-expressing cells (WT; n = 150 kinetochores from 30 cells and 3 replicates, *Hs*KI; n = 150 kinetochores from 30 cells and 3 replicates). (E) Representative images of BubR1 (magenta) in cells expressing WT Spc105 or the FUS-*Hs*KI chimera (green). (F) Quantification of BubR1 intensity relative to EGFP signal in WT and FUS-*Hs*KI-expressing cells (WT; n = 150 kinetochores from 30 cells and 3 replicates, FUS-*Hs*KI; n = 150 kinetochores from 30 cells and 3 replicates). (G) Representative images of Rod (magenta) in cells expressing WT Spc105 or the FUS-*Hs*KI chimera (green). (H) Quantification of Rod intensity relative to EGFP signal in WT and FUS-*Hs*KI-expressing cells (WT; n = 155 kinetochores from 31 cells and 3 replicates, FUS-*Hs*KI; n = 160 kinetochores from 32 cells and 3 replicates). (I) Percentage of anaphase/telophase cells with at least one lagging chromosome in WT-, deletion mutant- and chimera-expressing cells (n = 150 cells per condition from 3 replicates). (J) Percentage of WT-, deletion mutant- and chimera-expressing cells with at least one lagging merotelic in anaphase or cut chromatin in telophase. (K) Model for how Spc105 and BubR1 contribute to recruitment of a core population of RZZ at bioriented KTs that reduces the Ndc80 complex’s affinity for kMTs. Dynein relieves the core RZZ population’s weakening of kMT attachment affinity via stripping in fly cells. Reported p-values were generated by a randomization method: ** p-value < 0.001. Boxed regions are shown in the insets. Error bars are SEM. Scale bars: 5 μm, 1μm (insets). Letters above each plot column indicate significant differences (p < 0.05) determined by two-tailed t-tests for all pairwise combinations. Same letters indicate that the difference is not significant (p > 0.05).

Altogether our findings support a model whereby the N-terminus of Spc105 possesses “sticky” IDRs that mediate localized higher-order multimerization of KI motifs to fully recruit BubR1 and RZZ to bioriented kinetochores. This phenomenon allows Spc105 to localize sufficient levels of a core pool of RZZ necessary to promote accurate chromosome segregation by reducing the frequency of merotelic attachments. The kinetochore is inherently multivalent in nature and there are 100s of Spc105/KNL1 molecules within the outer kinetochore region. Our findings indicate that the “baseline” level of multivalency at the kinetochore is insufficient to fully recruit RZZ. Rather, we propose that inter- and intra-molecular interactions of sticky IDRs in Spc105 supports an additional level of higher-order oligomerization in the outer kinetochore that increases the avidity of Spc105 for RZZ. The KI motifs in Spc105 are insufficient to support the functional linkage to RZZ and dynein since Spc105-mCherry-Cry2 fusions that contained the KI motifs never exhibited polar accumulation/localization. However, the linkage was established on a seconds timescale once higher-order multimerization was triggered as evidenced by immediate dynein-dependent motility of clusters upon photo-activation. It is presently unclear how many molecules of Spc105 must be present in the higher-order oligomers to support RZZ recruitment since the number of molecules in Cry2 oligomers is unclear. Future experiments with oligomerization tags of known stoichiometries will be informative to define the minimal number of Spc105 molecules that are necessary to support the functional linkage to RZZ and dynein. Nonetheless, we propose that assembly/enrichment of proteins into the multivalent kinetochore structure may not always be sufficient for their functionality. In these instances, enrichment may stimulate localized higher-order oligomerization within the structure to support emergent properties of the kinetochore.

Our data are entirely consistent with a prior model of the mechano-regulatory crosstalk pathway in which RZZ decreases the affinity of the Ndc80 complex (core binding factor) for MTs until it is inhibited by dynein via stripping or conformational change (Amin et al., 2018; Cheerambathur et al., 2013). While earlier work envisioned that a coronal pool of RZZ functions early in mitosis to prevent premature end-on attachments, we propose that kMT attachment stability is also actively being modulated by a core pool of RZZ throughout mitosis - even at bioriented kinetochores. Our findings also further flesh out details of the molecular model for the crosstalk pathway by identifying Spc105 as a receptor for the kMT attachment-regulating pool of RZZ. More specifically, we propose that recruitment of the core pool of RZZ is coordinated by a complex of Spc105 bound to BubR1, which binds to the KI2 motif in Spc105 via its N-terminal TPR domain (**Fig. 5 K**). In *D. melanogaster (Dm)* cells, dynein strips RZZ, but not BubR1, suggesting that the affinity between BubR1 and Spc105 is stronger than RZZ’s affinity for BubR1 and Spc105. We cannot rule out that there are additional direct contacts between the N-terminal IDRs in Spc105 and RZZ as Rod was reduced but not absent from kinetochores in cells expressing the Δ266-384 and in BubR1 depleted cells. We also cannot exclude that Bub1 contributes to RZZ recruitment in flies as it was moderately enriched in the Spc105 clusters. The N-terminus of KNL1 and Bub1 are required for full recruitment of RZZ to unattached kinetochores in human cells (Caldas et al., 2015) and BUB-1 is required for the localization of a pool of RZZ that recruits the dynein-dynactin complex to kinetochores in *C. elegans* (Edwards et al., 2018). Interestingly, a region in *Dm*BubR1 possesses significant homology to a Bub1 motif required for RZZ recruitment to human kinetochores (Zhang et al., 2019; Zhang et al., 2015). Thus, we postulate that the linkage to RZZ described here is likely mediated by KNL1 and Bub1 (rather than BubR1) outside flies (**Fig. S3B C**). It will be worthwhile to further dissect how the core pool of RZZ is recruited to kinetochores given its importance to accurate chromosome segregation. Furthermore, physically linking Spc105 to dynein may result in transient force transduction through the disordered protein that could affect its SAC-regulating function and may explain the previously described “unraveling” of human KNL1 that was proposed to act as a possible tension sensing mechanism (Roscioli et al., 2020).

## MATERIALS AND METHODS

### Cell culture and cell line production

*Drosophila* S2 cells were cultured at 24°C in Schneider’s medium supplemented with 10% heat-inactivated FBS (Invitrogen) and 0.5x antibiotic–antimycotic cocktail (Invitrogen). All cell lines were made by transfecting WT S2 cells with 1 – 2 μg of the appropriate plasmid DNA. The cells were transfected using the Effectene Transfection Reagent (Qiagen) according to the manufacturer’s protocol. Stable cell lines were generated by selection with the appropriate antibiotic (0.025 mg/ml Blasticidin S HCl (ThermoFisher) or 0.25 - 0.5 mg/ml Hygromycin B (Invitrogen)) until cell death ceased. Stable cells were occasionally split in the presence of antibiotic to maintain expression levels.

### Live-cell imaging

Cells were seeded onto Concanavalin A- (ConA) (Sigma-Aldrich) treated 35 mm glass bottom petri dishes (Cellvis) were imaged on a TiE inverted microscope (Nikon) equipped with a Borealis (Andor) retrofitted CSU-10 (Yokogawa) spinning disk head and ORCA-Flash4.0 LT Digital CMOS camera (Hamamatsu) using a 100x 1.49 numerical aperture (NA) Apo differential interference contrast (DIC) TIRF objective (Nikon). Metamorph software (Molecular Devices) was used to control the imaging system and to analyze data.

### Optogenetic photo-activation experiments and analysis in mitotic cells

The same imaging protocol was used to analyze the behavior of 1-400-mCherry-Cry2, 266-384-mCherry-Cry2, and 1-1722-mCherry-Cry2 in mitotic cells. Cells were induced ∼16 hours overnight with 500 μM CuSO_4_ and seeded onto ConA-coated coverslips before subjecting them to the activation/imaging protocol outlined in Fig. 1C, which consisted of a pre-activation image with 561 nm light, followed by blue light activation at 488 nm and synchronous imaging of 561 nm at 10 second intervals for 3 minutes.

For phenotypic assessment of the DHC knockdown experiments, the above protocol was used to produce confocal time-lapses of which the final image was analyzed by linescan to quantify the distribution of 1-1722-mCherry-Cry2 clusters along the length of the spindle. The line width was set to ∼30% of the spindle width and extended lengthwise along the long spindle axis terminating at each pole. Average fluorescence intensity values for each point along the line were transferred to a spreadsheet. Linescan data was pooled after normalizing for the variability in spindle length relative to the shortest spindle in a dataset. To do this, a running average of adjacent values was created for all spindles, and points were removed according to the extent to which a spindle needed to be normalized. For example, if a spindle needed to be corrected by 30%, every third data point was removed, but still accounted for by averaging adjacent data points along the linescan. For phenotypic assessment of the Rod knockdown experiments, linescans (drawn as described previously) were conducted on maximum projections of confocal Z-stacks of the mCherry channel following completion of the photo-activation protocol.

### Optogenetic clustering assays

*Drosophila* S2 cells stably expressing inducible mCherry-Cry2, 1-1722-mCherry-Cry2, or 266-384-mCherry-Cry2 were induced with 500 μM CuSO4 ∼16 hours prior to seeding them onto ConA-treated glass petri dishes. Since Spc105-B possesses MT binding activity, all chambers were treated with 25 μM colchicine for 2 hours prior to activation/imaging and then subjected to imaging for a maximum of 90 minutes at room temperature. A pre-activation image (561 nm excitation) was taken prior to the first photo-activation event (488 nm excitation) while acquiring a three-minute time-lapse with mCherry images taken every 15 seconds. All replicates paired a control (mCherry-Cry2) dish and an experimental dish (either 1-1722-mCherry-Cry2 or 266-384-mCherry-Cry2) each of which were subjected to identical activation/imaging protocols on the same day. To assess the expression levels necessary to support oligomerization, the average mCherry intensity of a 10×10 pixel region prior to photo-activation was recorded and plotted for every cell that assembled discernible clusters during the activation/imaging protocol.

### Fold enrichment assay

*Drosophila* S2 cells stably expressing inducible 266-384-mCherry-Cry2 were transiently transfected with plasmids encoding EGFP, EGFP-Bub1, Bub1-EGFP, Zw10-EGFP, sfGFP-Rod, or BubR1-EGFP each under low/endogenous-expression promoters using Effectene (Qiagen) according to the manufacturer’s protocol. Within 1 week of transfection, the cell lines were induced with 500 μM CuSO4 for ∼16 hours prior to seeding onto ConA-treated glass petri dishes and subjecting them to the activation/imaging protocol outlined in Fig. 1C. A confocal Z-section was then taken of each field at the conclusion of the 3-minute photo-activation protocol. The time-lapses were first assessed to determine if there was evident co-localization (obvious increase by eye of EGFP signal above the local background in the vicinity of 266-384 droplets) and the average EGFP intensity within a 10-pixel X 10-pixel region placed in a representative region of the cytoplasm was recorded from the pre-activation image. This analysis was used to record the co-localization propensity as a function of expression levels of the co-transfected EGFP-tagged proteins shown in Fig. 2. Only cells exhibiting evident co-localization were subjected to the fold-enrichment analysis with the exception of the control EGFP-expressing cells, which never exhibited co-localization but were analyzed as described below to establish a baseline for the assay and for comparison purposes. To measure the fold enrichment of 266-384-mCherry-Cry2 binding partners, color combined planes from the confocal Z-sections were scanned to identify co-localized puncta. A 10-pixel X 10-pixel region was then centered on 266-384 droplets and transferred to the EGFP channel for analysis. The fold enrichment was defined as the integrated intensity of EGFP in the droplet divided by the integrated intensity of EGFP signal in the same region positioned nearby in cytoplasm lacking clusters. The fold enrichment of five droplets were measured per cell typically from different regions of the cell and from multiple planes.

### 1,6-hexanediole (HD) wash-in experiments and live-cell quantifications

To assess the effects of 1,6-HD on 266-384-mCherry-Cry2 drop assembly, cells were subjected to a 1 second photoactivation with 488 nm light but shortly after drops had assembled the media in the chamber was replaced with media supplemented with 3.5% 1,6-HD and time-lapse imaging of the mCherry channel continued. To assess the localization and levels of active Aurora B kinase, two ConA-coated coverslips were seeded with wildtype S2 cells and after providing time for the cells to adhere/flatten (∼15 minutes) the coverslips were treated with 10 μM MG132 for 1 hour. The media on the coverslips was then removed and replaced with media containing either MG132 (control) or MG132 and 3.5% 1,6-HD. After 3 minutes, the coverslips were fixed, stained, and quantified as described below. Two assays were used to assess the effects of 1,6-HD on the localization of Spc105, BubR1, and RZZ components. To assess kinetics, confocal time-lapse imaging of metaphase cells expressing Spc105-EGFP, BubR1-EGFP, Zw10-EGFP, or sfGFP-Rod was conducted as the media in the chamber was replaced with media supplemented with 3.5% 1,6-HD. The background corrected EGFP fluorescence of the metaphase plate was then measured over time using the duplicate region method (Ye and Maresca, 2018), normalized to the time-point prior to introducing 1,6-HD. To better quantify the extent to which the localization of each protein was affected by 1,6-HD, chambers of cells expressing each of the tagged proteins were treated for 1 hour with 10 μM MG132 and the stage positions of multiple (11 – 31) metaphase cells were saved in the software. Each of the marked metaphase cells were then subjected to confocal Z-stack time-lapses where the first Z-stack was taken prior to adding 1,6-HD and the second Z-stack was taken 3 minutes after the media was replaced with media containing MG132 and 3.5% 1,6-HD. The background corrected EGFP fluorescence of the metaphase plate regions were then measured from the maximum projections before and after introducing 1,6-HD using the duplicate region method. The percent of EGFP signal remaining 3 minutes post-HD addition relative to the pre-HD time-point is reported.

### Immunofluorescence and quantification

For immunofluorescence, cells were seeded on ConA-coated coverslips and after ∼1 hour the coverslips were rinsed in 1X BRB80 and fixed in either 100% methanol at −20°C (DHC and CID) or 10% paraformaldehyde in 1X BRB80 (all other conditions) for 10 minutes. Cells were permeabilized in 1X PBS + 1% Triton X-100 for 8 minutes, rinsed 3X in in 1X PBS + 0.1% Triton X-100, and then blocked in 5% boiled donkey serum (Jackson Immunoresearch) for 45 minutes. Cells were then incubated in the appropriate primary antibodies diluted in boiled donkey serum for 1 hour. Next, the coverslips were washed 3X, five minutes each, with PBS + 0.1% Triton X-100 and then incubated in the appropriate secondary antibodies (Jackson Immunoresearch) diluted (1:200 - 1:500) in boiled donkey serum supplemented with 0.1 mg/ml DAPI for 45 minutes. The coverslips were washed 3X (5 minutes each) in PBS + 0.1% Triton X-100 and then mounted in mounting media solution containing 90% glycerol, 20mM Tris pH 8.0, and 0.5% N-propyl gallate. The following conditions and antibodies were used for immunofluorescence-based experiments. Cells were treated with 25 μM colchicine for 1 hour prior to fixation for DHC and Rod RNAi quantifications. Cells were treated for 1 hour with 10 μM MG132 for quantifying Rod, BubR1, and pABK at bioriented kinetochores/centromeres. Primary antibodies: mouse anti-Dynein Heavy Chain (Developmental Studies Hybridoma Bank) at 1:200, rabbit anti-CID (Abcam) at 1:200, rabbit anti-Rod serum (gift of M. Przewloka) at 1:500, rabbit anti-BubR1 serum (gift of C. Sunkel) at 1:500 - 1:1000, sheep anti-Spc105 serum (gift of M. Przewloka) at 1:500 – 1:1000, rabbit anti-Phospho-aurora A/B/C (Cell Signaling Technology) at 1:1000, chicken anti-GFP (Abcam) at 1:1000, rabbit anti-phospho-H3 (Abcam) at 1:5000, and mouse anti-DM1α (Sigma Aldrich) at 1:1000.

The region-in-region method (Ye and Maresca, 2018) was used to background correct and quantify integrated fluorescence intensities of the protein of interest ratioed to the indicated reference proteins at individual kinetochores/centromeres from confocal Z-planes for the following experiments: RNAi knockdowns of DHC and Rod, BubR1 and Rod kinetochore levels in Δ266-384- and Δ266-384 chimera-expressing cells, and kinetochore levels in BubR1-depleted cells. To measure centromeric levels of pABK in DMSO- and HD-treated cells a 10-pixel X 10-pixel box was centered between two sister kinetochores and the duplicate region method was used to measure the background corrected integrated fluorescence intensity, which was then ratioed to the sum of the background corrected fluorescence intensities of Spc105 signals from each sister kinetochore measured with the region-in-region method. To quantify the BubR1 RNAi depletion, the region-in-region method was used on maximum projections of confocal Z-sections. To assess chromosome segregation phenotypes, Spc105-EGFP, Δ266-384-Spc105-EGFP, Δ266-384-Spc105-*Hs*KI-GFP, and Δ266-384-Spc105-FUS-*Hs*KI-EGFP cells were seeded on ConA-coated coverslips, fixed, and stained for phospho-H3, GFP, and α-tubulin (DM1α). Cells in late anaphase were scored as containing a merotelic when at least one evident (often stretched) GFP-positive kinetochore was observed in between the segregating chromosomes. A cell was defined as having a bridge when at least one chromosome was positioned between the segregating chromosomes but without any evident GFP-positive kinetochores. Cells were scored as having cut chromatin when decondensing chromatin was stretched between the two masses of segregated chromosomes often resulting in the chromatin being pinched or cut by the cytokinetic furrow.

### DNA Constructs

All DNA constructs were made using isothermal (Gibson) cloning (Gibson et al., 2009) into a pMT-V5-His B vector. The mCherry-Cry2 constructs were made by amplifying mCherry-Cry2 DNA from Addgene plasmid #101221 (gift from Brangwynne). The PCR product was ligated, using a 5’ KpnI site and a 3’ ApaI site, downstream of the copper inducible metallothionine promoter. The 1-400 Spc105-mCherry-Cry2 construct was made by amplifying Spc105-B corresponding to amino acids 1-400, which was previously amplified from cDNA (DGRC Clone IP22012). This fragment was flanked by KpnI sites and inserted into the vector previously described, between the metallothionein promoter and the mCherry-Cry2. The 266-384-mCherry-Cry2 and 1-1722 Spc105-mCherry-Cry2 constructs were made by amplifying Spc105-B cDNA sequence corresponding to amino acids 266-384 (beginning with a start codon) and 1-1722, respectfully, with flanking KpnI sites. The DNA was inserted into the pMT vector (containing mCherry-Cry2) as previously described. The Bub1-EGFP construct was made by amplifying Bub1 from Bub1 cDNA (LD22858). The PCR product was inserted into a pMT vector downstream of a Cenp-C promoter and upstream of EGFP via a 5’ SpeI site and a 3’ XbaI site. The N-terminal EGFP-Bub1 construct was made by amplifying Bub1 from cDNA (LD22858), and inserting the product, flanked by SacII sites, at the 3’ end of GFP, downstream of a Cenp-C promoter. The BubR1-EGFP construct was made by amplifying BubR1 promoter and gene from genomic DNA and inserting it into the pMT vector upstream of EGFP via a 5’ XhoI site and a 3’ NotI site. The ZW10-EGFP construct was made by amplifying ZW10 from cDNA CG9900-RB. The DNA was inserted downstream of the KNL1 promoter and upstream of EGFP using a 5’ XhoI site and a 3’ XbaI site. sfGFP-Rod was a gift from Gohta Goshima. The Spc105 Δ266-384 construct was made by amplifying cDNA sequence corresponding to amino acids 1-265 and 385-1959 with overlapping primers engineered to position BamHI and BcuI sites between the two fragments for the purpose of generating the chimeras. The DNA pieces were inserted downstream of the KNL1 promoter and upstream of EGFP using a 5’ XhoI site and a 3’ XbaI site. The Spc105 *Hs*KI chimera construct was made by inserting the human KI motif region amplified from cDNA sequence corresponding to amino acids 201-249 into a BcuI site between 1-265 Spc105 and 385-1959 Spc105. The Spc105 FUS-HsKI chimera construct was made by inserting the human FUS disordered region in front of the KI motifs in the Spc105 Δ266-384 HsKI construct, flanked by a BamHI site.

### RNA Interference

Dynein template was made by amplifying ∼600 base pairs from the dynein heavy chain cDNA (CG7507-RA). The T7 promoter sequence (5′-TAATACGACTCACTATAGGG-3′) was followed by (5′-TGCCCAGGCGAATAGTTGGT-3′) in the forward primer and (5′- CAAGTTTAAAGTATTTCATT-3′) in the reverse. Rod template was made by amplifying ∼600 base pairs from an sfGFP-Rod construct (gift from Gohta Goshima). The T7 promoter sequence was followed by (5′-CGTCGCGAGGCATCTTCCAA-3′) in the forward primer and (5′- GTGCTTTGATCTCCAGTGAT-3′) in the reverse. The templates were then used to produce dsRNA using the T7 RiboMAX Express Large Scale RNA Production System (Promega) according to the manufacturer’s protocol. Stable cells expressing inducible 1-1722-mCherry-Cry2 were treated with 20ug of dsRNA products (DHC or Rod) or control RNA in serum-free media for 1 hour, followed by the addition of 1 mL of serum containing media and 2-(DHC) or 4-day (Rod) incubations at 24C before imaging.

### Statistical analyses

Statistical analyses reporting p-values from Student’s t-tests were done in excel while p-values generated with the randomization method were done using the PlotsofDifferences web tool at: https://huygens.science.uva.nl/PlotsOfDifferences (Goedhart, 2019). PlotsofDifferences calculates p-values using a randomization method that does not rely on any assumptions about the distribution of the data (normal versus non-normal).

## ACKNOWLEDGMENTS

We are grateful to Jennifer Le and Thomas Laskarzewski for help generating DNA constructs. Thank you to Marcin Przewloka (Rod and Spc105 sera), Gohta Goshima (sfGFP-Rod construct), and Claudio Sunkel (BubR1 serum) for reagents as well as to Kim McKim for discussing unpublished observations. pHR-mCh-Cry2WT and pHR-FUSN-mCh-Cry2WT were gifts from Clifford Brangwynne (Addgene plasmids #101221; http://n2t.net/addgene:101221; RRID:Addgene_101221 and #101223; http://n2t.net/addgene:101223; RRID:Addgene_101223). This work was supported by an NIH grant (GM107026) to T.J.M. and by an NIH T32 training grant that supported J.M.M. (GM135096) as a fellow in the UMass Biotechnology Training Program.

## SUPPLEMENTAL MATERIALS

### Supplemental Figures

**Figure S1.**
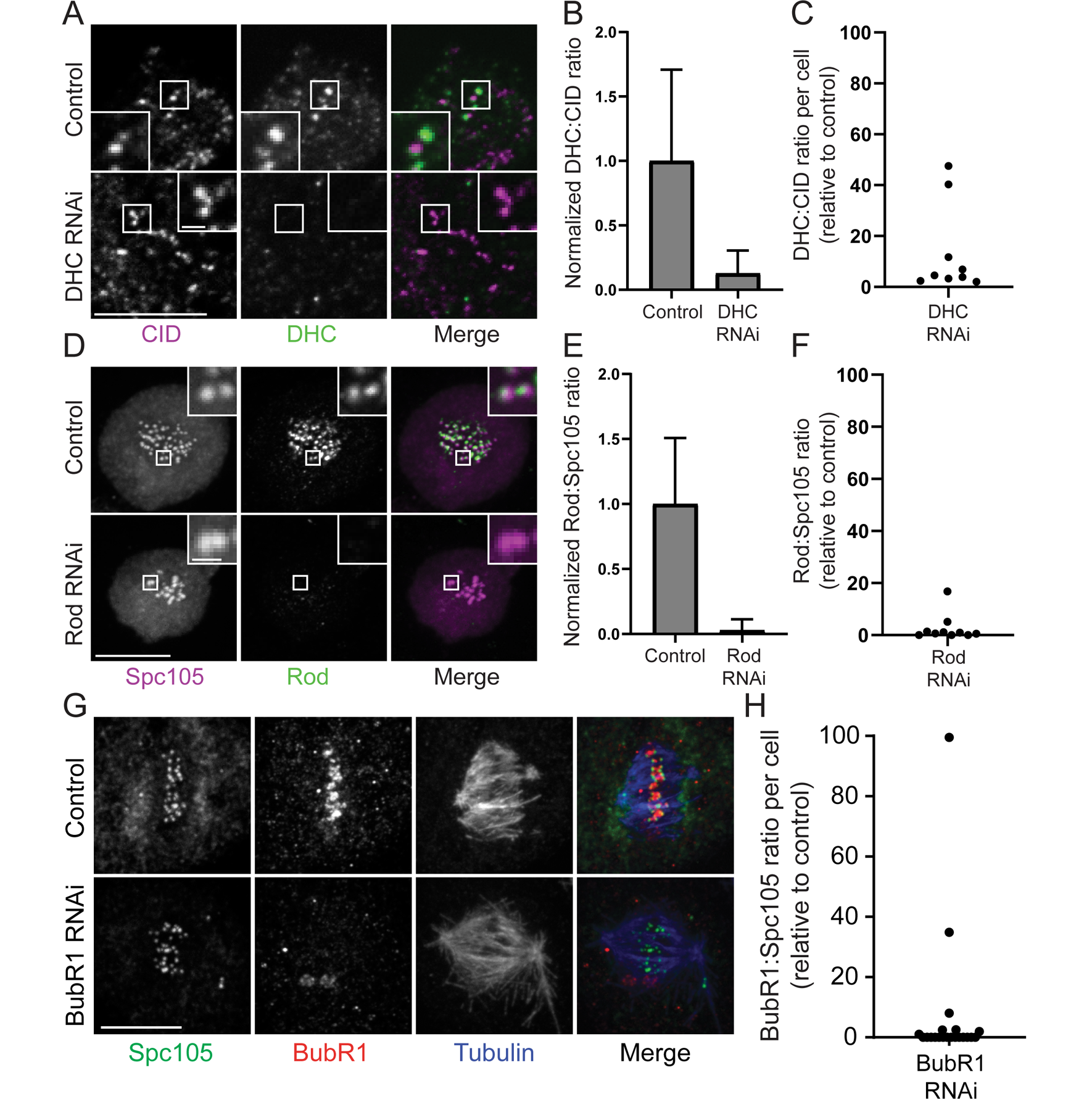
Characterization of DHC, Rod, and BubR1 depletions. (A) Representative confocal Z-planes of control and DHC RNAi cells stained for CID (magenta) and DHC (green). (B) Quantification of DHC ratioed to CID fluorescence (n = 50 kinetochores from 10 cells for each condition). (C) Cell averages (5 kinetochores quantified per cell) of DHC to CID ratios for DHC dsRNA-treated cells relative to the mean DHC:CID ratio of control cells as a measure of the DHC RNAi penetrance. (D) Representative confocal Z-planes of control and Rod RNAi cells stained for Spc105 (magenta) and Rod (green). (E) Quantification of Rod ratioed to Spc105 fluorescence (n = 50 kinetochores from 10 cells for each condition). (F) Cell averages (5 kinetochores quantified per cell) of Rod to Spc105 ratios for Rod dsRNA-treated cells relative to the mean Rod:Spc105 ratio of control cells as a measure of the Rod RNAi penetrance. (G) Representative maximum projections of confocal Z-sections of control and BubR1 dsRNA-treated fixed cells stained for Spc105 (green), BubR1 (red), and tubulin (blue). (H) Cell averages of BubR1:Spc105 ratios for BubR1 dsRNA-treated cells relative to the mean BubR1:Spc105 ratio of control cells as a measure of the BubR1 RNAi penetrance (control; n = 20 cells from 2 replicates, BubR1 RNAi; n = 21 cells from 2 replicates). Boxed regions are highlighted in the insets. Error bars are SD. Scale bars, 10 μm.

**Figure S2.**
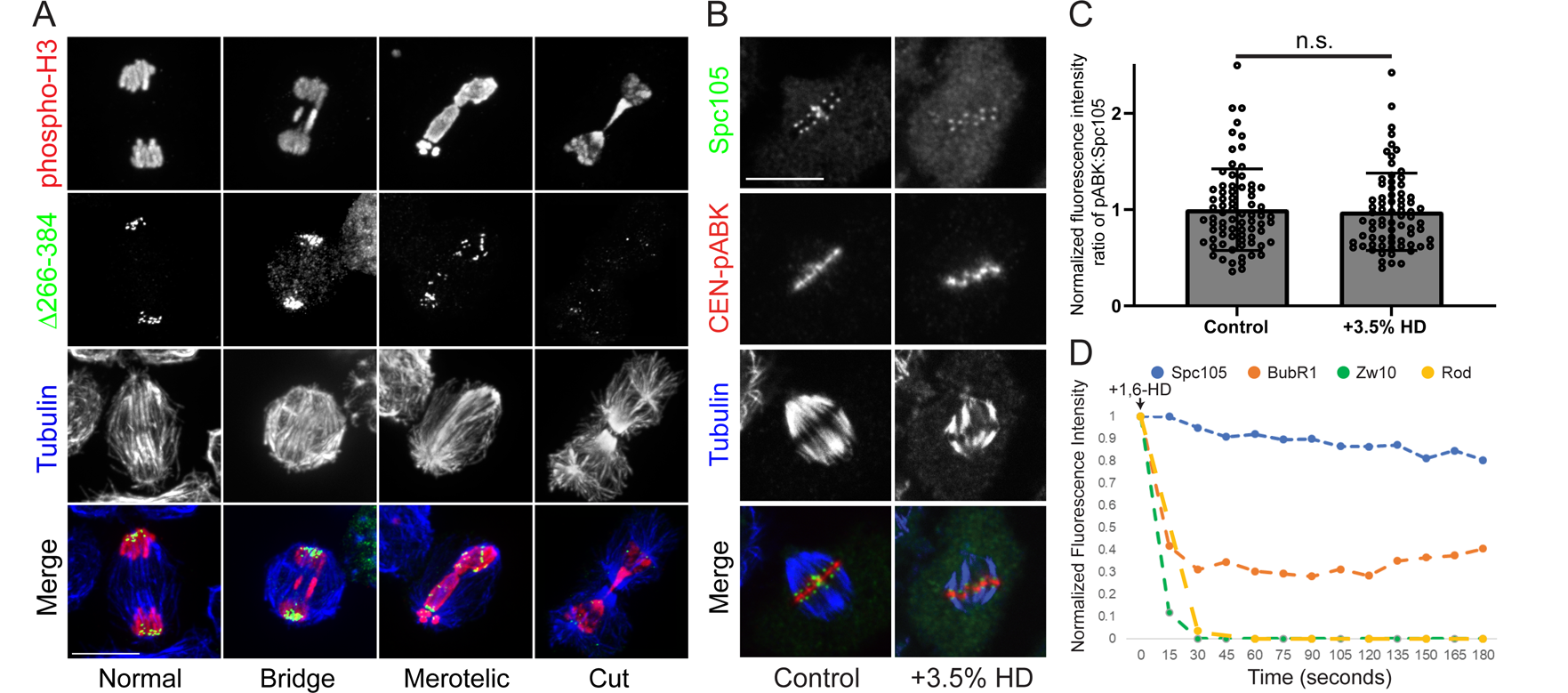
Chromosome mis-segregation examples in KI motif deletions and the assessing the effects of 1,6 HD on regulators of kMT attachment stability. (A) Chromosome segregation categories from Fig. 3G stained for phospho-H3 (red), EGFP (green), tubulin (blue). (B) Representative confocal Z-planes of cells fixed three minutes after washing in fresh media (control) or media supplemented with 3.5% 1,6-HD and stained for Spc105 (green), centromere-enriched phosphorylated (active) Aurora B kinase (CEN-pABK; red) and tubulin (blue). (C) Quantification of centromeric pABK signal ratioed to Spc105 signal (control; n = 75 kinetochore pairs/inner centromeres from 15 cells, HD; n = 73 kinetochore pairs/inner centromeres from 15 cells). (D) Quantification of the chromosome-proximal fluorescence intensities of the indicated EGFP-tagged proteins from the representative time-lapses shown in Fig. 4E. Error bars are SD. Scale bars, 10 μm.

**Figure S3.**
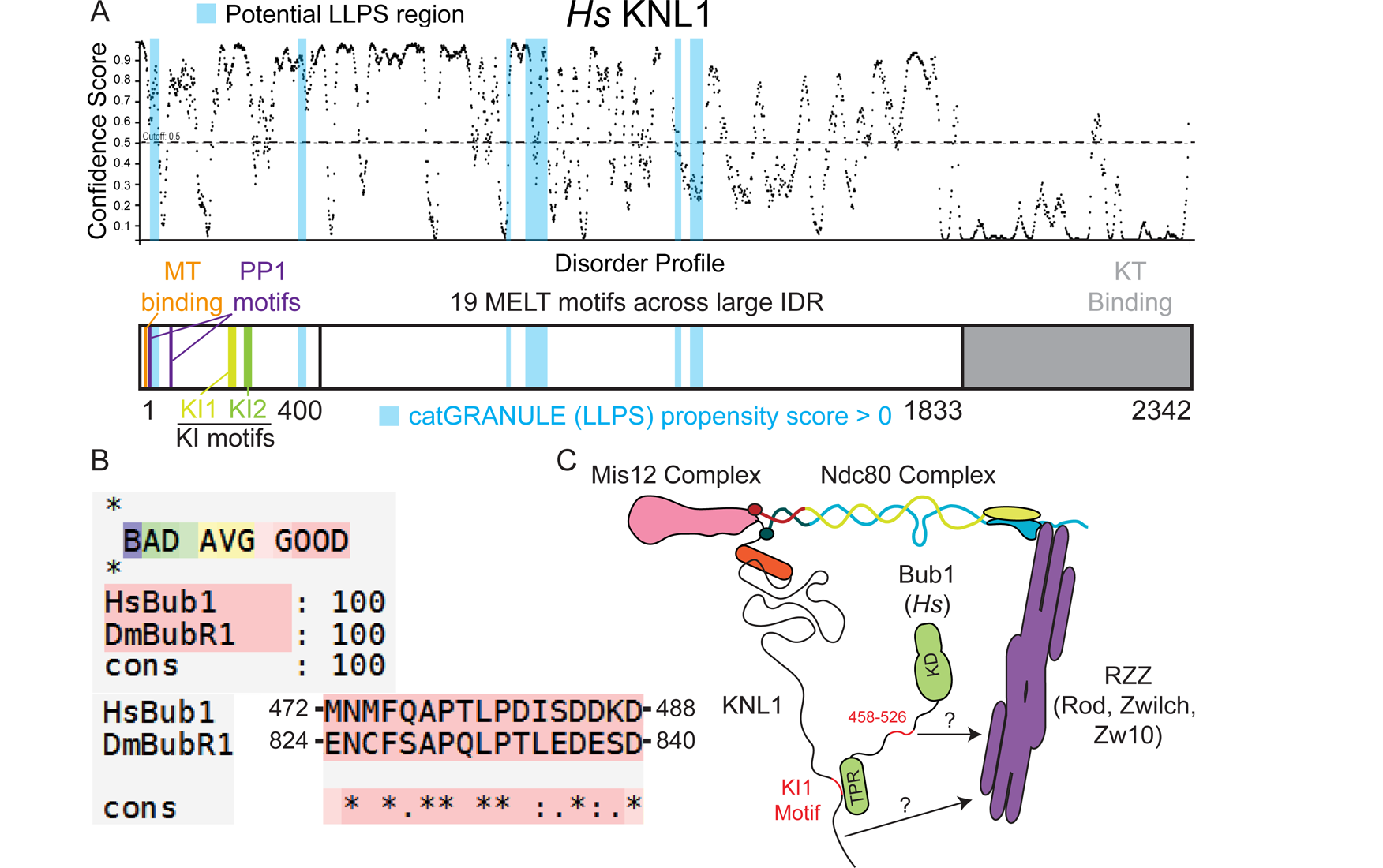
KI motifs and LLPS regions do not overlap in human KNL1 and KNL1/Bub1 may be an RZZ receptor in human cells. (A) Schematic of human (*Hs*) KNL1 highlighting the N-terminal MT-(orange) and PP1-binding (magenta) motifs, the KI motifs (shades of green), and the six regions across Spc105 with a catGRANULE LLPS propensity score above 0 (blue) overlaid onto a corresponding disorder plot of KNL1. (B) T-COFFEE sequence alignment of human (Hs) Bub1 and *D. melanogaster* (Dm) BubR1 amino acid sequences identifying a motif in *Drosophila* BubR1 with strong homology to a region in human Bub1 that contributes to recruiting RZZ to the kinetochore. (C) Speculative molecular model of how the pathway characterized using *D. melanogaster* cells in this study could function in human cells where the KNL1-based RZZ receptor may be Bub1 rather than BubR1.

## Supplemental Videos

**Video 1. Spinning disk confocal time-lapse of a mitotic cell expressing Spc105-1-400-mCherry-Cry2 during the photo-activation protocol**. Displayed times are min:sec. Scale bar, 10 μm. Relates to Fig. 1D.

**Video 2. Spinning disk confocal time-lapses of control (left) and DHC-depleted (right) mitotic cells expressing Spc105-1-1722-mCherry-Cry2 during the photo-activation protocol.** Displayed times are min:sec. Scale bar, 10 μm. Relates to Figs. 1E, F.

**Video 3. Spinning disk confocal time-lapses of control (left) and Rod-depleted (right) mitotic cells expressing Spc105-1-1722-mCherry-Cry2 during the photo-activation protocol.** Displayed times are min:sec. Scale bar, 10 μm. Relates to Figs. 1G, H.

**Video 4. Two-color spinning disk confocal time-lapse of Rod (green) localized to a 266-384-mCherry-Cry2 droplet (magenta) moving poleward.** Displayed times are min:sec. Scale bar, 10 μm. Relates to Fig. 2M.

**Video 5. Spinning disk confocal time-lapses of representative merotelic attachments in anaphase cells expressing the Spc105 Δ266-384 deletion mutant.** Displayed times are min:sec. Scale bars, 10 μm. Relates to Fig. 3J.

**Video 6. Spinning disk confocal time-lapses of cells expressing comparable levels of mCherry-Cry2 (upper left), 1-1722-mCherry-Cry2 (upper right), and 266-384-mCherry-Cry2 (lower left) during application of the same photo-activation protocol**. Displayed times are min:sec. Scale bar, 10 μm. Relates to Fig. 4A.

**Video 7. Spinning disk confocal time-lapse of a representative 266-384-mCherry-Cry2 oligomer fusion event**. Displayed times are min:sec. Scale bar, 1 μm. Relates to Fig. 4C.

**Video 8. Spinning disk confocal time-lapse of 266-384-mCherry-Cry2 oligomerization followed by wash-in of 1,6-HD and rapid cluster disassembly**. Displayed times are min:sec. Scale bar, 10 μm. Relates to Fig. 4D.

**Video 9. Spinning disk confocal time-lapses of representative 1,6-HD wash-in experiments in metaphase cells expressing Spc105-EGFP (upper left), BubR1-EGFP (upper right), Zw10-EGFP (lower left), and sfGFP-Rod (lower right)**. Displayed times are min:sec. Scale bar, 10 μm. Relates to Figs. 4E and S2B.

